# findPC: An R package to automatically select number of principal components in single-cell analysis

**DOI:** 10.1101/2021.10.15.464460

**Authors:** Haotian Zhuang, Zhicheng Ji

## Abstract

**Summary:** Principal component analysis (PCA) is widely used in analyzing single-cell genomic data. Selecting the optimal number of PCs is a crucial step for downstream analyses. The elbow method is most commonly used for this task, but it requires one to visually inspect the elbow plot and manually choose the elbow point. To address this limitation, we developed six methods to automatically select the optimal number of PCs based on the elbow method. We evaluated the performance of these methods on real single-cell RNA-seq data from multiple human and mouse tissues. The perpendicular line method with 20 PCs has the best overall performance, and its results are highly consistent with the numbers of PCs identified manually. We implemented the six methods in an R package, `findPC`, that objectively selects the number of PCs and can be easily incorporated into any automatic analysis pipeline.

**Availability and Implementation:** findPC R package is freely available at https://github.com/haotian-zhuang/findPC

**Supplementary information:** Supplementary data are available at *Bioinformatics* online.

## 1 Introductions

Principal Component Analysis (PCA) is a fundamental dimension reduction technique in analyzing single-cell genomic data. It maps the cells with high-dimensional and noisy genomic information to a low-dimensional and denoised principal component space. The principal components (PCs) are then used to group cells into clusters [1], identify continuous cell trajectories [2], or serve as the basis of other dimension reduction techniques such as t-SNE [3] and UMAP [4]. The number of PCs plays a critical role in downstream analyses. With too many PCs, PCs with the smallest variations may represent the noise in the data and dilute the signal. With too few PCs, the essential information in the data may not be captured. An optimal number of PCs should keep the essential information in the data while filtering out as much noise as possible.

The elbow method is most commonly used to determine the number of PCs in single-cell analysis literature. It starts from a scatterplot where the y-axis shows the standard deviations of PCs and the x-axis shows the numbers of PCs. Since the PCs are ordered decreasingly by their standard deviations, the scatterplot usually presents a monotonically decreasing curve that sharply descends for the first several PCs and gradually flattens out for the subsequent PCs. The optimal number of PCs is then visually identified as the elbow point where the curve bends from steep to flat descent. However, visual identification of elbow points requires manual work and cannot be incorporated into an automatic pipeline. Plus, the results may not be reproduced by different researchers for PCs with ambiguous elbow points.

To address these issues, we propose six methods to automatically identify the elbow point, thus the number of PCs in single-cell analysis. We applied these methods to real single-cell RNA-seq data from multiple human or mouse tissues and evaluated their performance using the elbow points manually annotated by us. The results show that the method based on perpendicular lines with 20 PCs has the best overall performance. We implemented these functions in a user-friendly R package, findPC, that is freely available on Github.

## 2 Methods

We developed six methods to computationally identify elbow points based on different heuristics (Figure 1A-1F). The details of these methods are included in Supplementary Materials. The input to these methods is a vector of standard deviations of the first *N* PCs with the largest standard deviations. *N* has different choices of 20, 30, 40, 50.

**Fig. 1.**
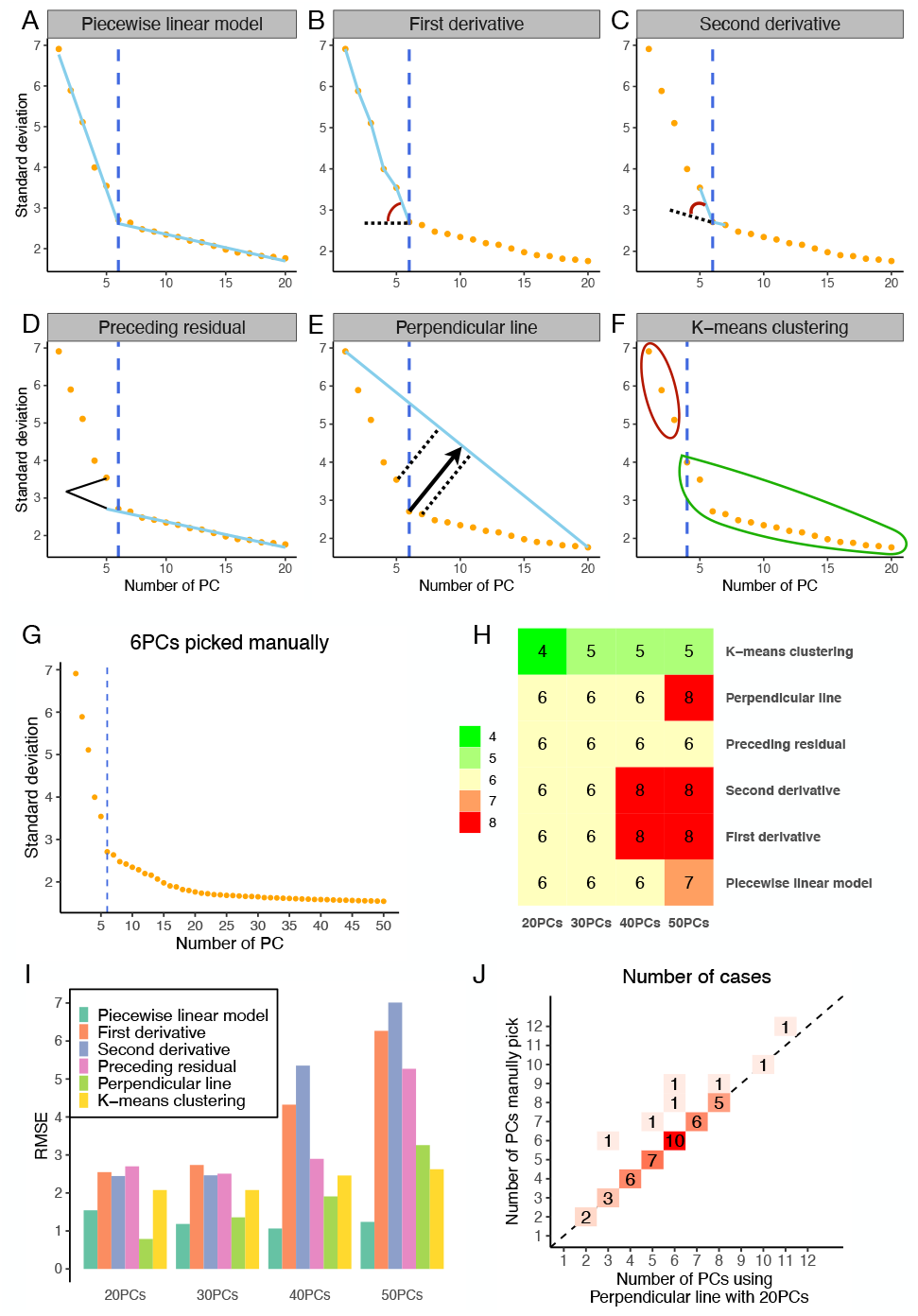
A-F. Demonstration of the six methods that automatically identify the optimal number of PCs. The elbow plot is obtained using the human fetal brain tissue. The top 20 PCs are shown. The blue dashed vertical line shows the elbow point chosen by each method. G. Elbow plot for the human fetal brain tissue. The top 50 PCs are shown. The visually identified elbow point (6 PCs) is shown as the blue dashed vertical line. H. Number of PCs chosen by six methods using different total numbers of PCs (*N*) for the human fetal brain tissue. I. Overall performance of six methods using different total numbers of PCs (*N*), measured by RMSE across all selected tissues. J. Number of PCs chosen manually (y-axis) or by perpendicular line with 20 PCs (x-axis). Numbers in the colored squares represent the numbers of tissues.

Method 1 fits a series of continuous piecewise linear models with two pieces to the scatterplot and chooses the PC with the best fit (Figure 1A). Method 2 chooses the last PC whose absolute value of the first derivative on the elbow plot is larger than a cutoff learned from data (Figure 1B). Method 3 is similar to Method 2, except it depends on the second derivatives (Figure 1C). Method 4 fits a linear regression line using the last few PCs and calculates the residual of the preceding PC. It iterates through all possible PCs and chooses the last PC with preceding residual larger than a cutoff learned from data (Figure 1D). Method 5 chooses the PC with the longest perpendicular line to the line passing the two points of the first and last PCs on the elbow plot (Figure 1E). Method 6 groups all PCs into two clusters based on their standard deviations and chooses the first PC in the cluster with a smaller averaged standard deviation (Figure 1F).

To evaluate the performance of the six methods, we collected single-cell RNA-seq data of 65 human tissues from Human Cell Landscape [5] and 54 mouse tissues from Mouse Cell Atlas [6]. For each tissue, we used standard Seurat pipeline [1] to process the data, perform PCA analysis, and obtain the standard deviation of each PC. We then visually inspected the elbow plot for each tissue, selected 25 human tissues and 21 mouse tissues with clear elbow points, and identified the position of the elbow point for each tissue.

## 3 Results

For each selected human or mouse tissue, elbow points were automatically chosen by the six methods and compared to the visually identified elbow point. Figure 1G shows an example elbow plot with visually identified elbow point (6 PCs) in human fetal brain tissue, the same tissue used in Figure 1A-1F. Figure 1H shows the elbow points identified by different methods and using different total numbers of PCs (*N*). Elbow points identified by most methods agree with the visually identified elbow point in this specific tissue. The elbow plots and visually or automatically identified elbow points for all selected tissues are included in Supplementary Figure 1 for human tissues and Supplementary Figure 2 for mouse tissues. Figure 1I shows the overall performance of different methods, where root-mean-square errors (RMSEs) are calculated between visually identified elbow points and automatically chosen elbow points across all human and mouse tissues for each method. Method 5 (Perpendicular line) with top 20 PCs has the best overall performance, with RMSE 0.79. In almost all tissues, elbow points identified by Method 5 with top 20 PCs either agree with or are very close to elbow points identified visually (Figure 1J). Supplementary Figure 3 shows the comparisons of visually and automatically identified elbow points for all six methods and with different total numbers of PCs (*N*).

## 4 Implementations

We developed an R package, `findPC`, that implements the six methods. The R package has one function, `findPC`, that accepts a numeric vector of standard deviations of PCs as input and by default returns an integer value of the automatically selected elbow point as output. Based on the evaluations, we choose Method 5 (Perpendicular line) with top 20 PCs as the default method.

The following R command runs `findPC` with the default settings: > findPC (sdev = sdev)

This function returns an integer value of number of PCs selected by Method 5 (Perpendicular line) with top 20 PCs. It can be directly fed into downstream analyses or incorporated in an automatic pipeline for analyzing single-cell genomic data. Users can also run `findPC` for one or multiple methods with different total numbers of PCs (*N*), or get a comprehensive list of results for all combinations of six methods and total numbers of PCs. The `findPC` function also supports the aggregation of results from multiple methods and total numbers of PCs by taking the average, the median, or the mode. For users who want to visually inspect the results, `findPC` can also display the elbow plot with the automatically identified elbow points. All these functionalities are described in detail in the package vignette of `findPC`.

## Acknowledgements

Z.J. was supported by the Whitehead Scholars Program at Duke University School of Medicine.

## References

[1] Tim Stuart, Andrew Butler, Paul Hoffman, Christoph Hafemeister, Efthymia Papalexi, William M Mauck III, Yuhan Hao, Marlon Stoeckius, Peter Smibert, and Rahul Satija. Comprehensive integration of single-cell data. Cell., 177(7):1888–1902, 2019.

[2] Zhicheng Ji and Hongkai Ji. Tscan: Pseudo-time reconstruction and evaluation in single-cell rna-seq analysis. Nucleic Acids Res., 44(13):e117–e117, 2016.

[3] Laurens Van der Maaten and Geoffrey Hinton. Visualizing data using t-sne. J Mach Learn Res., 9(11), 2008.

[4] Etienne Becht, Leland McInnes, John Healy, Charles-Antoine Dutertre, Immanuel WH Kwok, Lai Guan Ng, Florent Ginhoux, and Evan W Newell. Dimensionality reduction for visualizing single-cell data using umap. Nat Biotechnol., 37(1):38–44, 2019.

[5] Xiaoping Han, Ziming Zhou, Lijiang Fei, Huiyu Sun, Renying Wang, Yao Chen, Haide Chen, Jingjing Wang, Huanna Tang, Wenhao Ge, et al. Construction of a human cell landscape at single-cell level. Nature., 581(7808):303–309, 2020.

[6] Xiaoping Han, Renying Wang, Yincong Zhou, Lijiang Fei, Huiyu Sun, Shujing Lai, Assieh Saadatpour, Ziming Zhou, Haide Chen, Fang Ye, et al. Mapping the mouse cell atlas by microwell-seq. Cell., 172(5):1091–1107, 2018.

