## Supplementary Materials for "findPC: An R package to automatically select number of principal components in single-cell analysis"

### Mathematical formulation

Let $s_i$ be the standard deviation of the $i$th PC, where $i = 1, 2, ..., N$. Since PCs are ordered by their standard deviations in a decreasing order, we have $s_{i+1} < s_i$ ($i = 1, 2, ..., N-1$).

#### Piecewise linear model

A continuous piecewise linear model with two pieces joining at $p$ can be written as $s_i = f(i) + \epsilon$. Here $\epsilon$ is the random noise, and

$$f(i) = \begin{cases} \alpha_0 + \alpha_1 * i & \text{if } i \leq p \\ \beta_0 + \beta_1 * i & \text{if } i > p \end{cases}$$
$$s.t.\ \alpha_0 + \alpha_1 * p = \beta_0 + \beta_1 * p \tag{1}$$

For each $p$, findPC finds the least square fit of the model and calculates the sum of squared error (SSE): $\sum_{i=1}^{N}[s_i - f(i)]^2$. The optimal number of PCs is chosen as $p$ with the smallest SSE.

#### First derivative

We define $d_i$, the first derivative of $i$th PC, as $s_i - s_{i-1}$ ($i = 2, 3, ..., N$). A density curve is fitted to the distribution of $|d_i|$ using the `density` function in R. A cut-off $z$ is determined as the trough of the first two peaks in the density curve, or equivalently, the first location where the first derivative is zero and the second derivative is positive. The optimal number of PCs is chosen as the last PC with $|d_i|$ larger than $z$, or equivalently, $\arg\max_i |d_i| > z$.

#### Second derivative

We define $e_i$, the second derivative of $i$th PC, as $d_{i+1} - d_i = (s_{i+1} - s_i) - (s_i - s_{i-1})$ ($i = 2, 3, ..., N-1$). Similar to the first derivative method, a density curve is fitted to $e_i$, a cutoff is determined as the trough of the first two peaks, and the optimal number of PCs is chosen as the last PC with $e_i$ larger than the cutoff.

#### Preceding residual

For $i$th PC, a simple linear regression model is fitted where response variable is $s_j$ and independent variable is $j$ ($j = i, i+1, ..., N$). Let $\hat{s}_{i-1}$ be the predicted value for $(i-1)$th PC in the fitted regression model ($i = 2, 3, ..., N-1$). We define $r_i = s_{i-1} - \hat{s}_{i-1}$ as the preceding residual. Similar to the first and second derivative approaches, a density curve is fitted to $r_i$, a cutoff is determined as the trough of the first two peaks, and the optimal number of PCs is chosen as the last PC with $r_i$ larger than the cutoff.

#### Perpendicular line

Let $l_i$ be the length of the perpendicular line from point $(i, s_i)$ to the line passing $(1, s_1)$ and $(N, s_N)$. The optimal number of PCs is chosen as the PC with the maximum $l_i$.

#### K-means clustering

K-means clustering is used to group all PCs into two clusters based on $s_i$. The optimal number of PCs is chosen as the smallest PC in the cluster with a smaller averaged standard deviation.
