## Supplementary Figure 1 for "findPC: An R package to automatically select number of principal components in single-cell analysis"

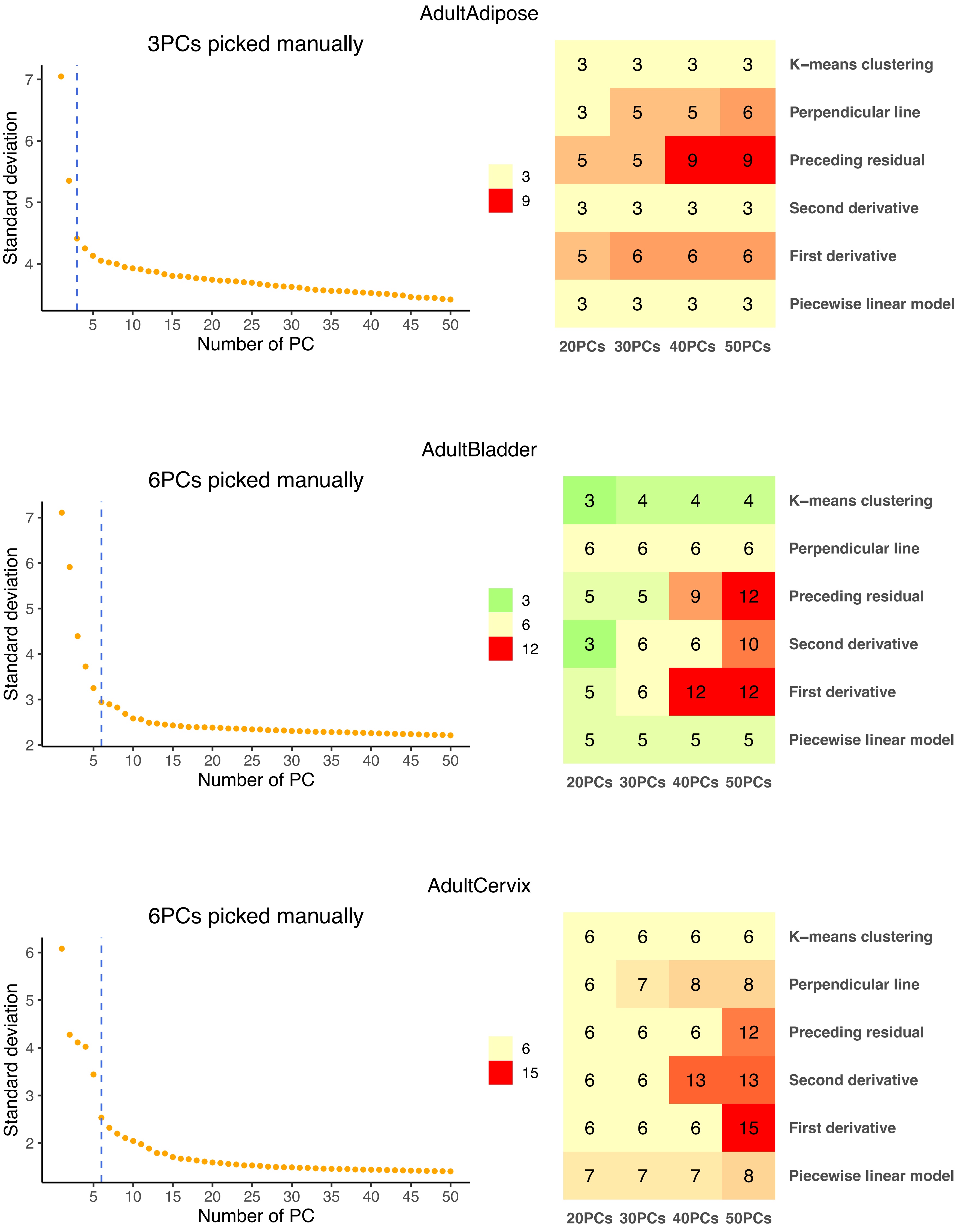
AdultAdipose
3PCs picked manually
Standard deviation
Number of PC
20PCs 30PCs 40PCs 50PCs
K−means clustering 3 3 3 3
Perpendicular line 3 5 5 6
Preceding residual 5 5 9 9
Second derivative 3 3 3 3
First derivative 5 6 6 6
Piecewise linear model 3 3 3 3
AdultBladder
6PCs picked manually
Standard deviation
Number of PC
20PCs 30PCs 40PCs 50PCs
K−means clustering 3 4 4 4
Perpendicular line 6 6 6 6
Preceding residual 5 5 9 12
Second derivative 3 6 6 10
First derivative 5 6 12 12
Piecewise linear model 5 5 5 5
AdultCervix
6PCs picked manually
Standard deviation
Number of PC
20PCs 30PCs 40PCs 50PCs
K−means clustering 6 6 6 6
Perpendicular line 6 7 8 8
Preceding residual 6 6 6 12
Second derivative 6 6 13 13
First derivative 6 6 6 15
Piecewise linear model 7 7 7 8

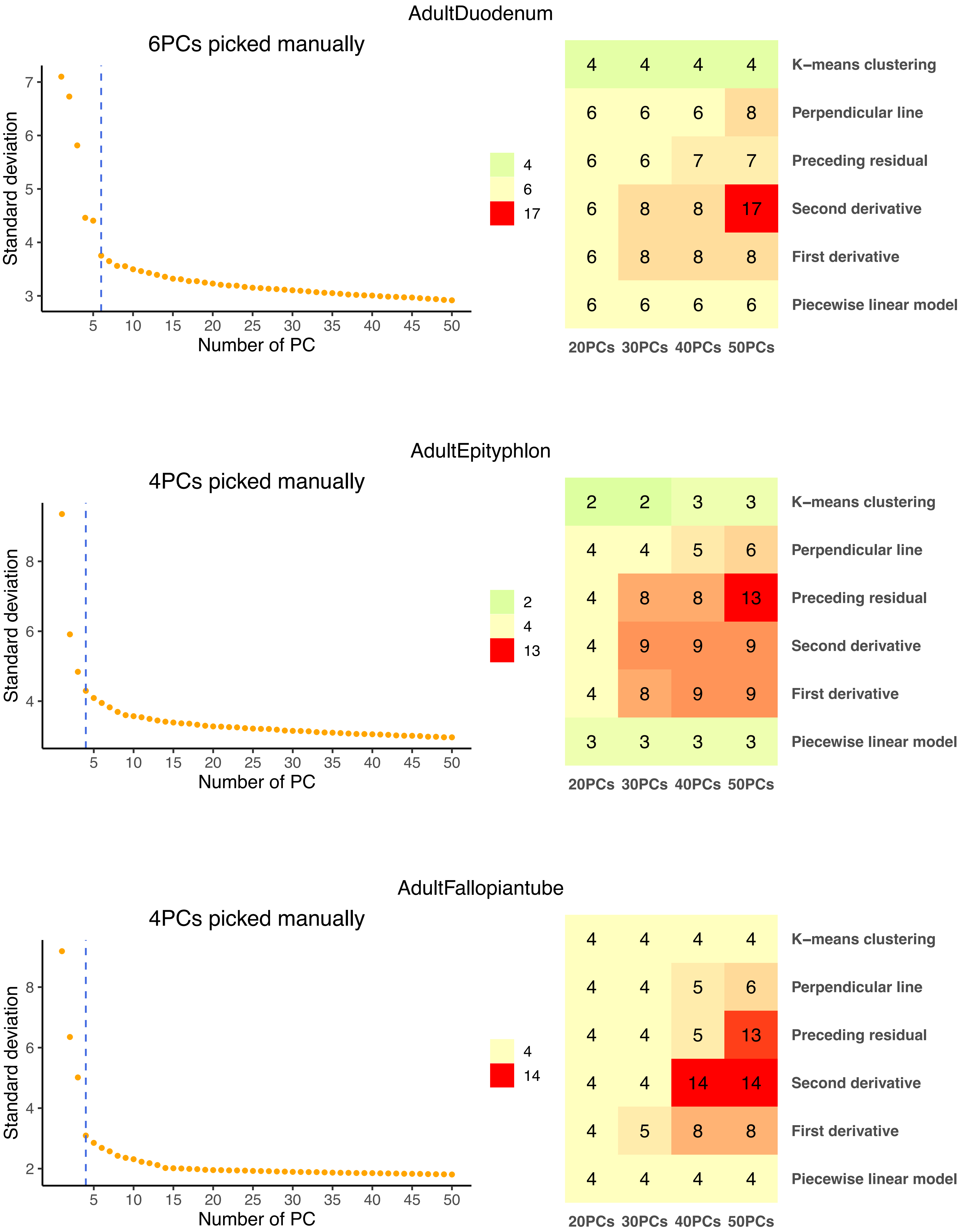
AdultDuodenum
6PCs picked manually
Standard deviation
Number of PC
K−means clustering
Perpendicular line
Preceding residual
Second derivative
First derivative
Piecewise linear model
20PCs 30PCs 40PCs 50PCs
4 4 4 4
6 6 6 8
6 6 7 7
6 8 8 17
6 8 8 8
6 6 6 6
AdultEpityphlon
4PCs picked manually
Standard deviation
Number of PC
K−means clustering
Perpendicular line
Preceding residual
Second derivative
First derivative
Piecewise linear model
20PCs 30PCs 40PCs 50PCs
2 2 3 3
4 4 5 6
4 8 8 13
4 9 9 9
4 8 9 9
3 3 3 3
AdultFallopiantube
4PCs picked manually
Standard deviation
Number of PC
K−means clustering
Perpendicular line
Preceding residual
Second derivative
First derivative
Piecewise linear model
20PCs 30PCs 40PCs 50PCs
4 4 4 4
4 4 5 6
4 4 5 13
4 4 14 14
4 5 8 8
4 4 4 4

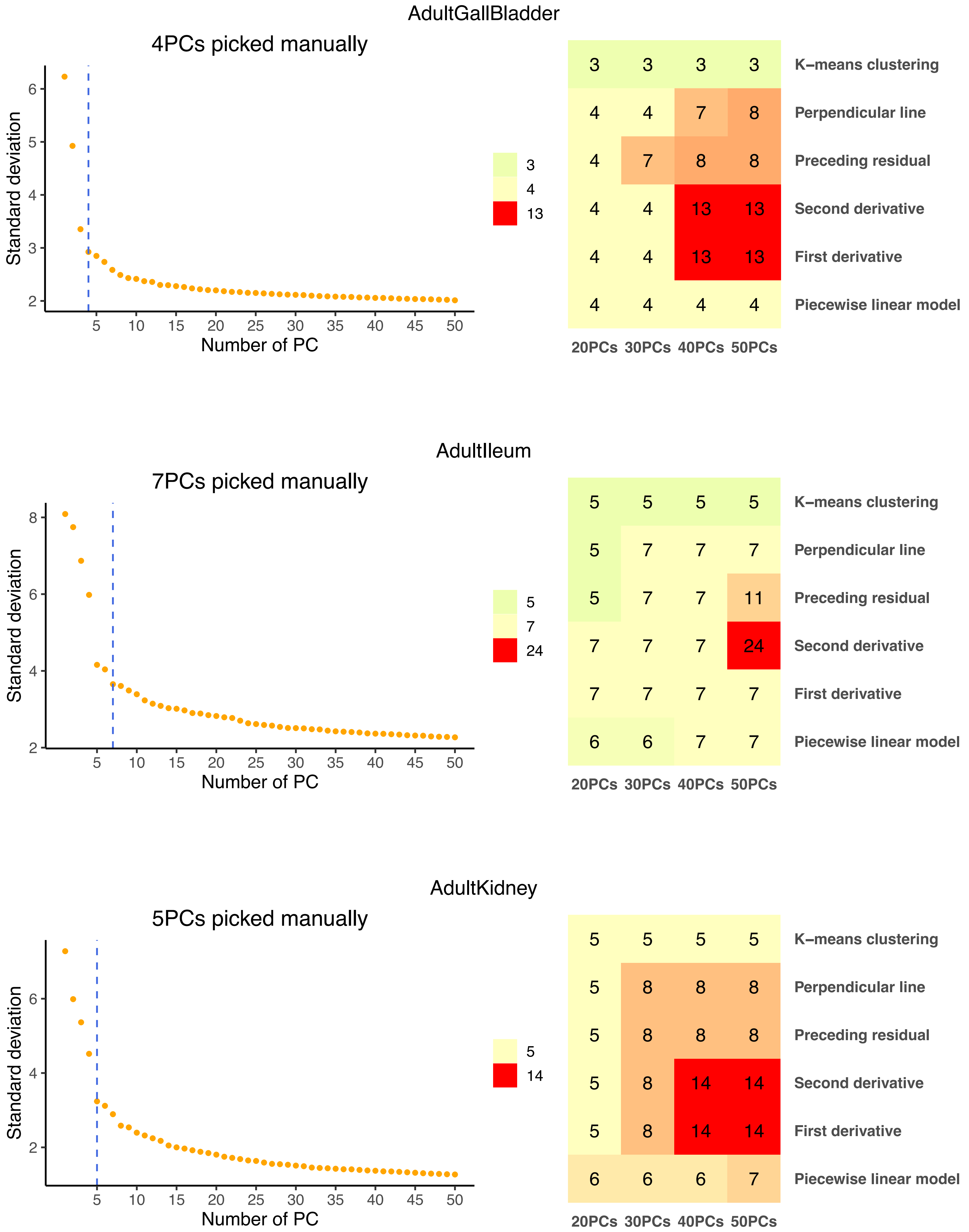
AdultGallBladder
4PCs picked manually
Standard deviation
Number of PC
3 3 3 3 K−means clustering
4 4 7 8 Perpendicular line
4 7 8 8 Preceding residual
4 4 13 13 Second derivative
4 4 13 13 First derivative
4 4 4 4 Piecewise linear model
20PCs 30PCs 40PCs 50PCs
3
4
13
AdultIleum
7PCs picked manually
Standard deviation
Number of PC
5 5 5 5 K−means clustering
5 7 7 7 Perpendicular line
5 7 7 11 Preceding residual
7 7 7 24 Second derivative
7 7 7 7 First derivative
6 6 7 7 Piecewise linear model
20PCs 30PCs 40PCs 50PCs
5
7
24
AdultKidney
5PCs picked manually
Standard deviation
Number of PC
5 5 5 5 K−means clustering
5 8 8 8 Perpendicular line
5 8 8 8 Preceding residual
5 8 14 14 Second derivative
5 8 14 14 First derivative
6 6 6 7 Piecewise linear model
20PCs 30PCs 40PCs 50PCs
5
14

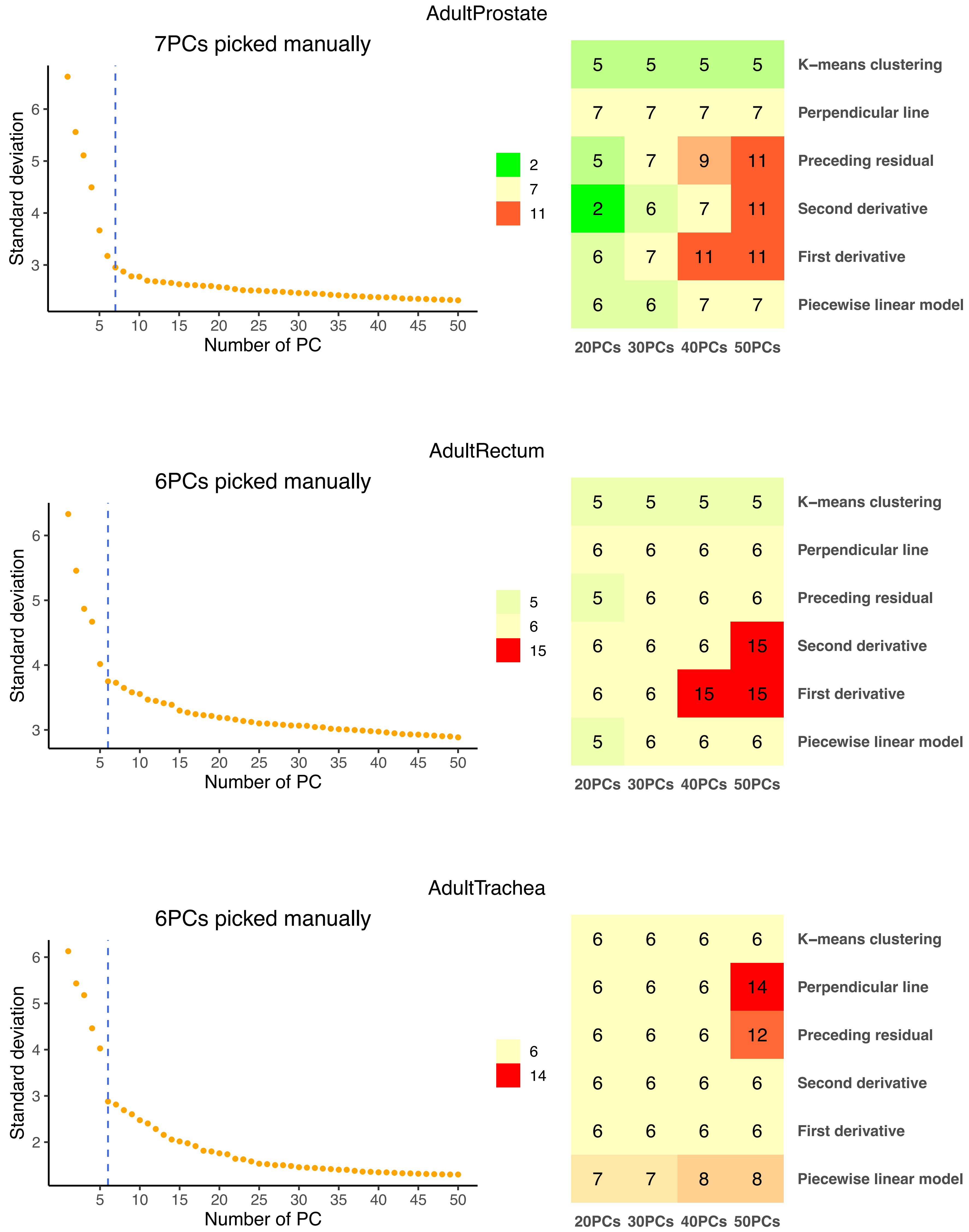
AdultProstate
7PCs picked manually
Standard deviation
Number of PC
2
7
11
5 5 5 5 K−means clustering
7 7 7 7 Perpendicular line
5 7 9 11 Preceding residual
2 6 7 11 Second derivative
6 7 11 11 First derivative
6 6 7 7 Piecewise linear model
20PCs 30PCs 40PCs 50PCs
AdultRectum
6PCs picked manually
Standard deviation
Number of PC
5
6
15
5 5 5 5 K−means clustering
6 6 6 6 Perpendicular line
5 6 6 6 Preceding residual
6 6 6 15 Second derivative
6 6 15 15 First derivative
5 6 6 6 Piecewise linear model
20PCs 30PCs 40PCs 50PCs
AdultTrachea
6PCs picked manually
Standard deviation
Number of PC
6
14
6 6 6 6 K−means clustering
6 6 6 14 Perpendicular line
6 6 6 12 Preceding residual
6 6 6 6 Second derivative
6 6 6 6 First derivative
7 7 8 8 Piecewise linear model
20PCs 30PCs 40PCs 50PCs

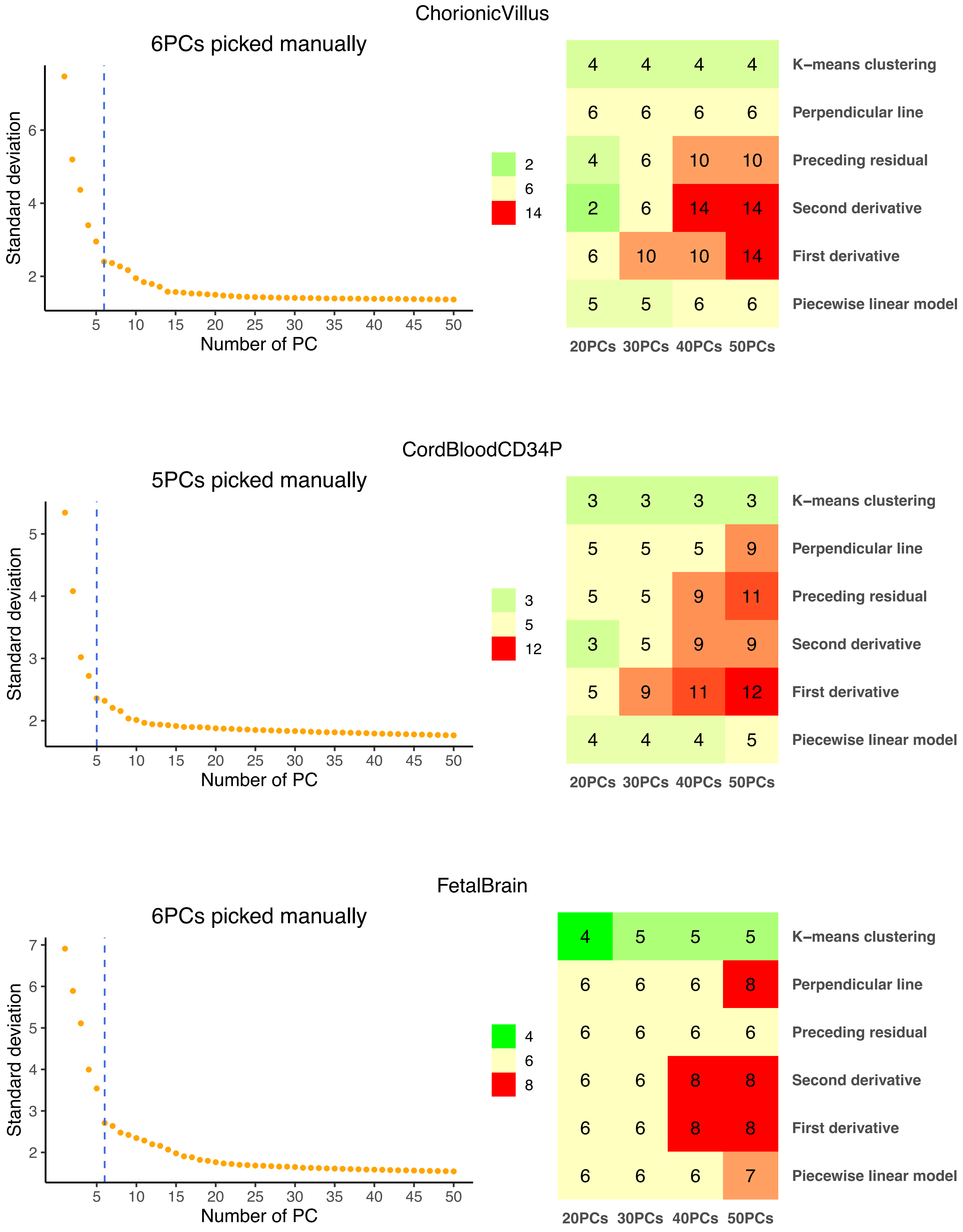
ChorionicVillus
6PCs picked manually
Standard deviation
Number of PC
2
6
14
K−means clustering: 4 4 4 4
Perpendicular line: 6 6 6 6
Preceding residual: 4 6 10 10
Second derivative: 2 6 14 14
First derivative: 6 10 10 14
Piecewise linear model: 5 5 6 6
20PCs 30PCs 40PCs 50PCs
CordBloodCD34P
5PCs picked manually
Standard deviation
Number of PC
3
5
12
K−means clustering: 3 3 3 3
Perpendicular line: 5 5 5 9
Preceding residual: 5 5 9 11
Second derivative: 3 5 9 9
First derivative: 5 9 11 12
Piecewise linear model: 4 4 4 5
20PCs 30PCs 40PCs 50PCs
FetalBrain
6PCs picked manually
Standard deviation
Number of PC
4
6
8
K−means clustering: 4 5 5 5
Perpendicular line: 6 6 6 8
Preceding residual: 6 6 6 6
Second derivative: 6 6 8 8
First derivative: 6 6 8 8
Piecewise linear model: 6 6 6 7
20PCs 30PCs 40PCs 50PCs

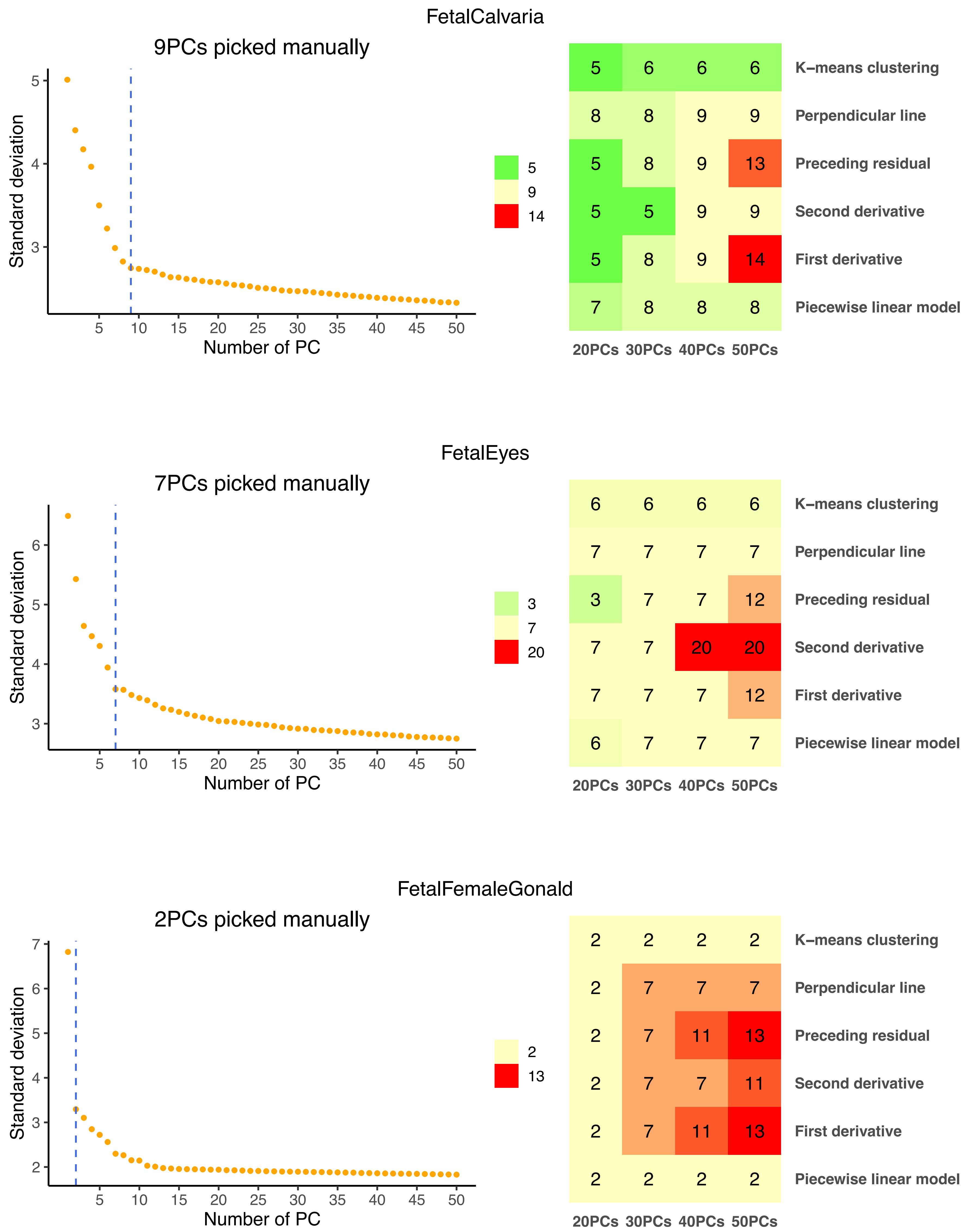
FetalCalvaria
9PCs picked manually
Standard deviation
Number of PC
K−means clustering 5 6 6 6
Perpendicular line 8 8 9 9
Preceding residual 5 8 9 13
Second derivative 5 5 9 9
First derivative 5 8 9 14
Piecewise linear model 7 8 8 8
20PCs 30PCs 40PCs 50PCs
FetalEyes
7PCs picked manually
Standard deviation
Number of PC
K−means clustering 6 6 6 6
Perpendicular line 7 7 7 7
Preceding residual 3 7 7 12
Second derivative 7 7 20 20
First derivative 7 7 7 12
Piecewise linear model 6 7 7 7
20PCs 30PCs 40PCs 50PCs
FetalFemaleGonald
2PCs picked manually
Standard deviation
Number of PC
K−means clustering 2 2 2 2
Perpendicular line 2 7 7 7
Preceding residual 2 7 11 13
Second derivative 2 7 7 11
First derivative 2 7 11 13
Piecewise linear model 2 2 2 2
20PCs 30PCs 40PCs 50PCs

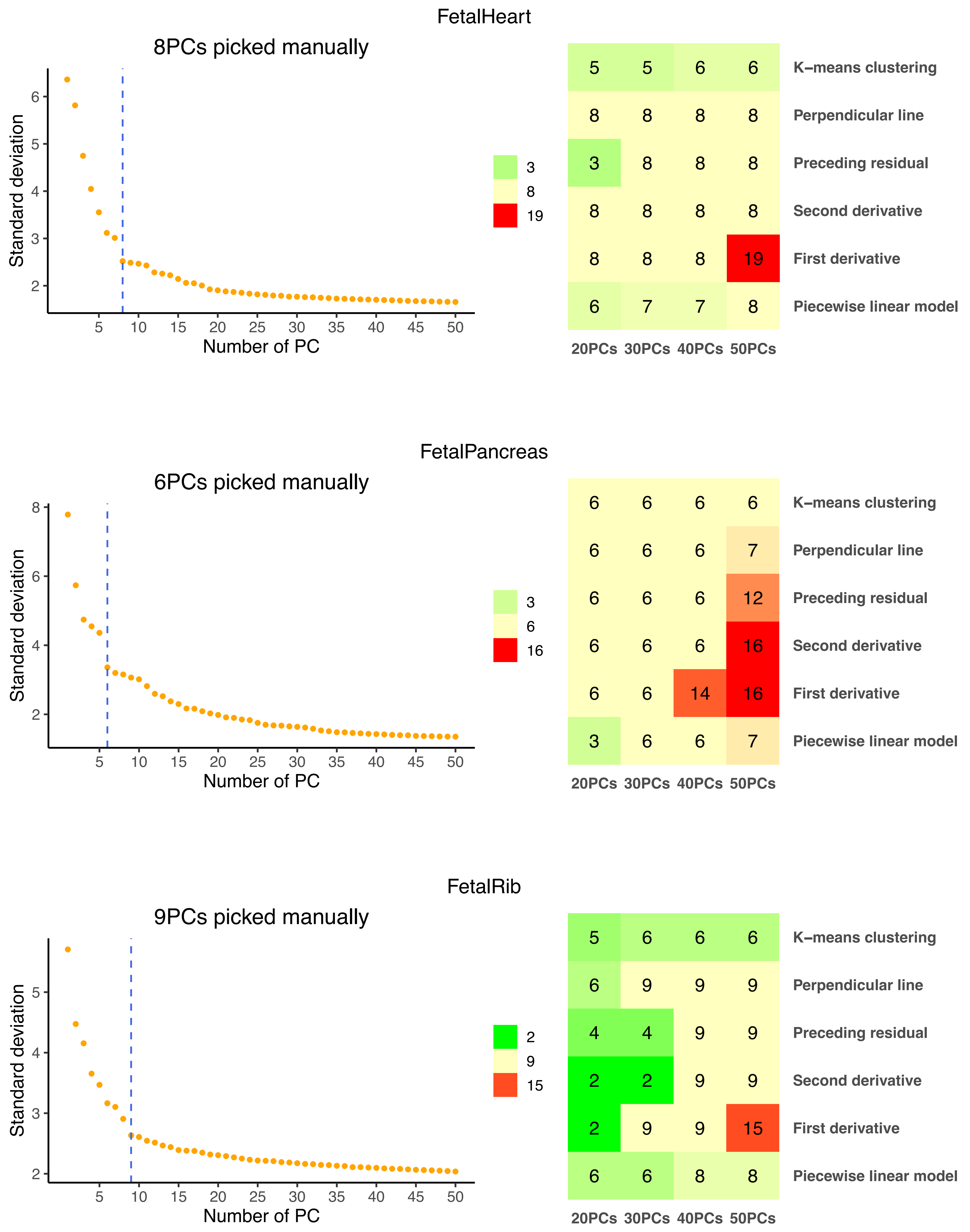
FetalHeart
8PCs picked manually
Standard deviation
Number of PC
K−means clustering
Perpendicular line
Preceding residual
Second derivative
First derivative
Piecewise linear model
20PCs 30PCs 40PCs 50PCs
FetalPancreas
6PCs picked manually
FetalRib
9PCs picked manually

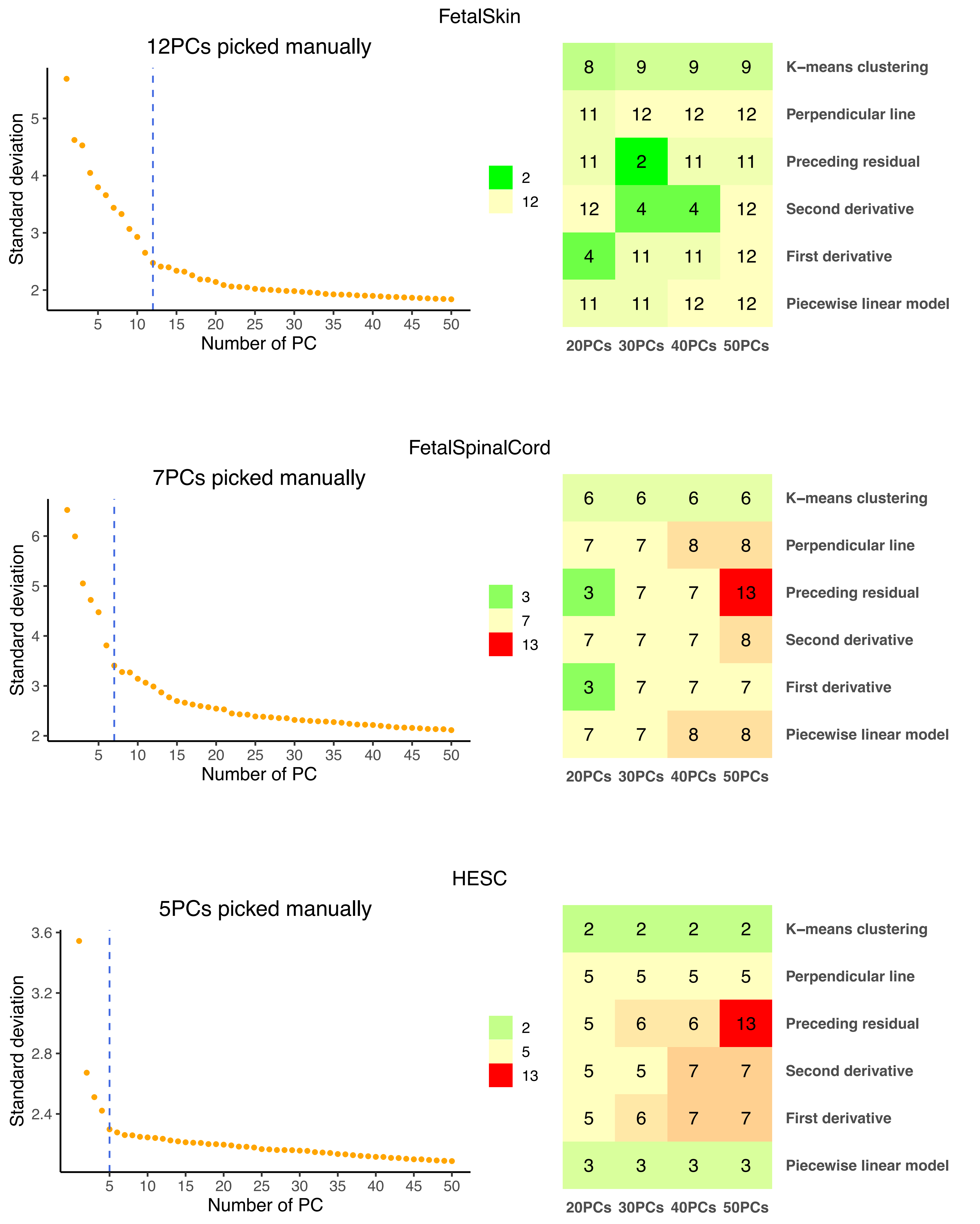
FetalSkin
12PCs picked manually
Standard deviation
Number of PC
5
10
15
20
25
30
35
40
45
50
2
3
4
5
2
12
8 9 9 9
11 12 12 12
11 2 11 11
12 4 4 12
4 11 11 12
11 11 12 12
K−means clustering
Perpendicular line
Preceding residual
Second derivative
First derivative
Piecewise linear model
20PCs 30PCs 40PCs 50PCs
FetalSpinalCord
7PCs picked manually
Standard deviation
Number of PC
2
3
4
5
6
3
7
13
6 6 6 6
7 7 8 8
3 7 7 13
7 7 7 8
3 7 7 7
7 7 8 8
K−means clustering
Perpendicular line
Preceding residual
Second derivative
First derivative
Piecewise linear model
20PCs 30PCs 40PCs 50PCs
HESC
5PCs picked manually
Standard deviation
Number of PC
2.4
2.8
3.2
3.6
2
5
13
2 2 2 2
5 5 5 5
5 6 6 13
5 5 7 7
5 6 7 7
3 3 3 3
K−means clustering
Perpendicular line
Preceding residual
Second derivative
First derivative
Piecewise linear model
20PCs 30PCs 40PCs 50PCs

Liver

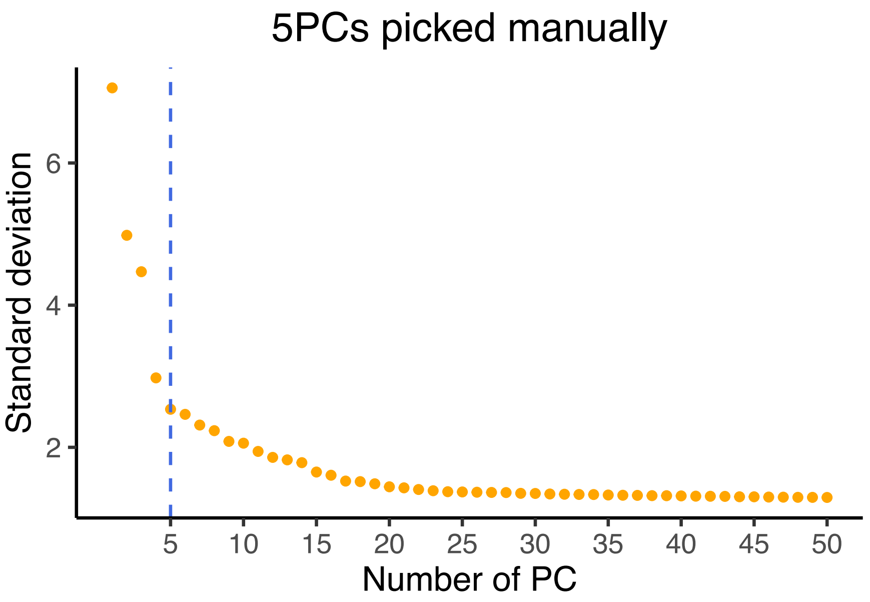


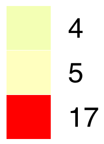


| | 20PCs | 30PCs | 40PCs | 50PCs |
|---|---|---|---|---|
| **K−means clustering** | 4 | 4 | 4 | 4 |
| **Perpendicular line** | 5 | 5 | 5 | 5 |
| **Preceding residual** | 5 | 5 | 5 | 5 |
| **Second derivative** | 4 | 5 | 5 | 5 |
| **First derivative** | 5 | 5 | 5 | 17 |
| **Piecewise linear model** | 4 | 5 | 5 | 5 |
