## Supplementary figures and images for "findPC: An R package to automatically select number of principal components in single-cell analysis"

### Supplementary Figure 2

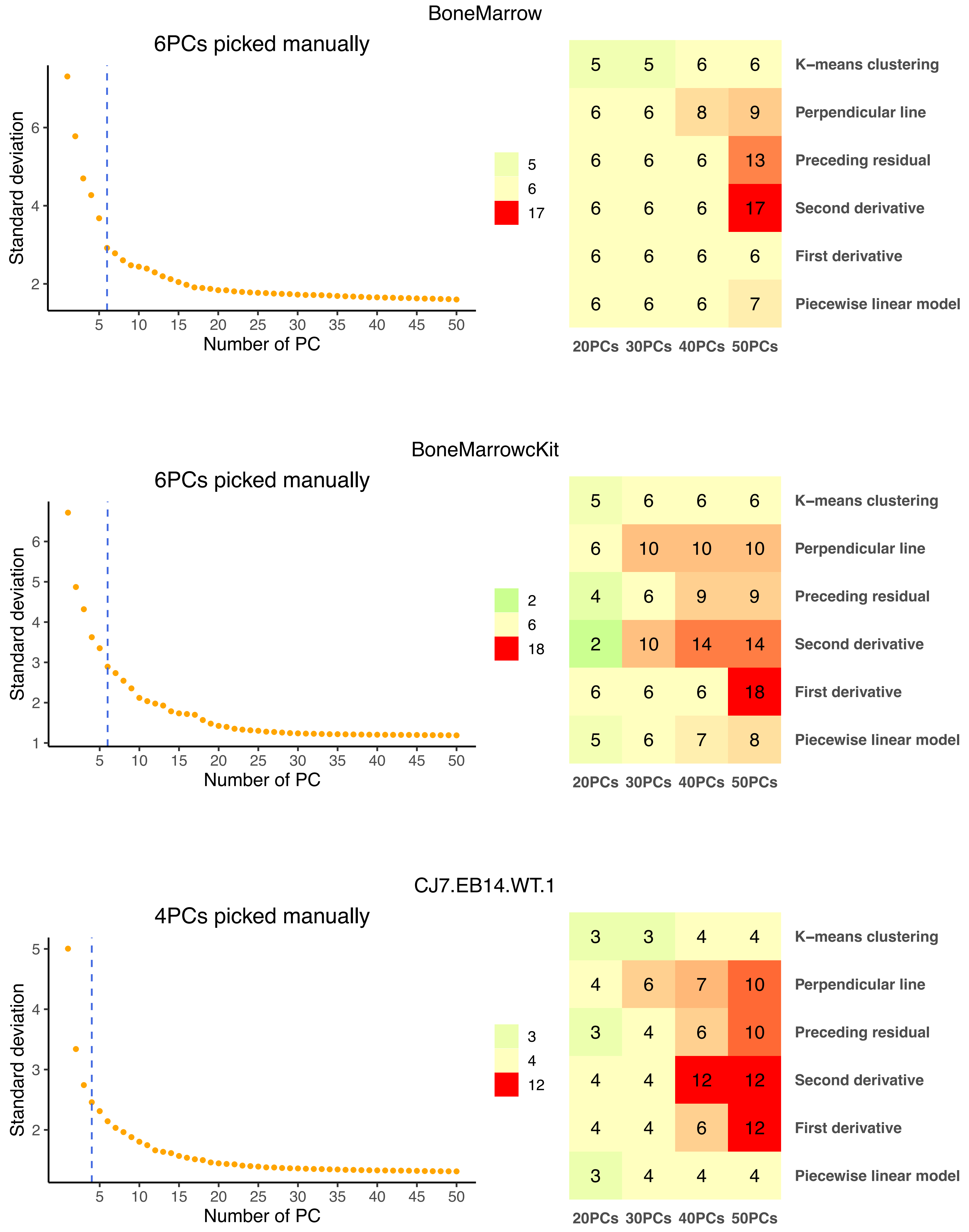

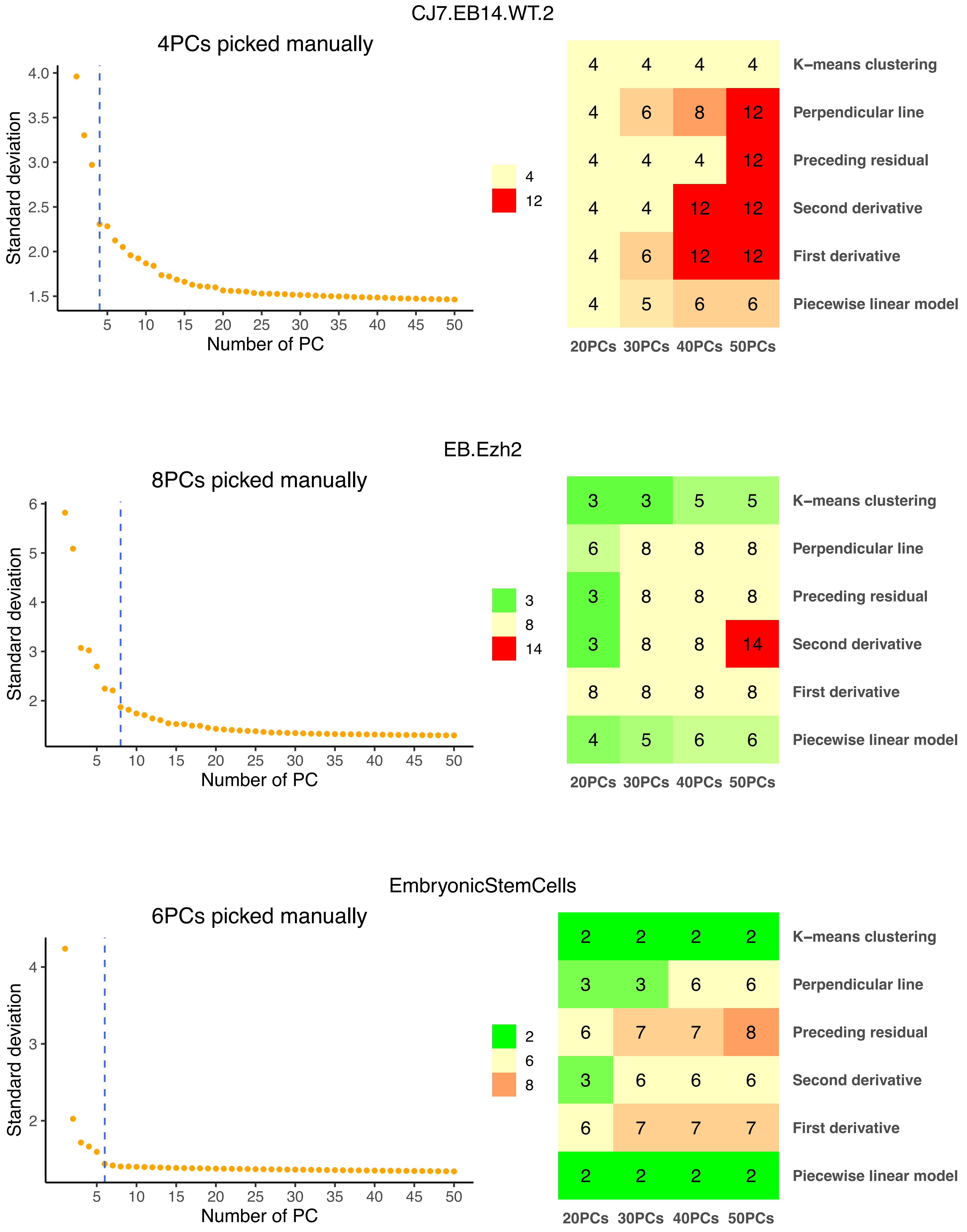

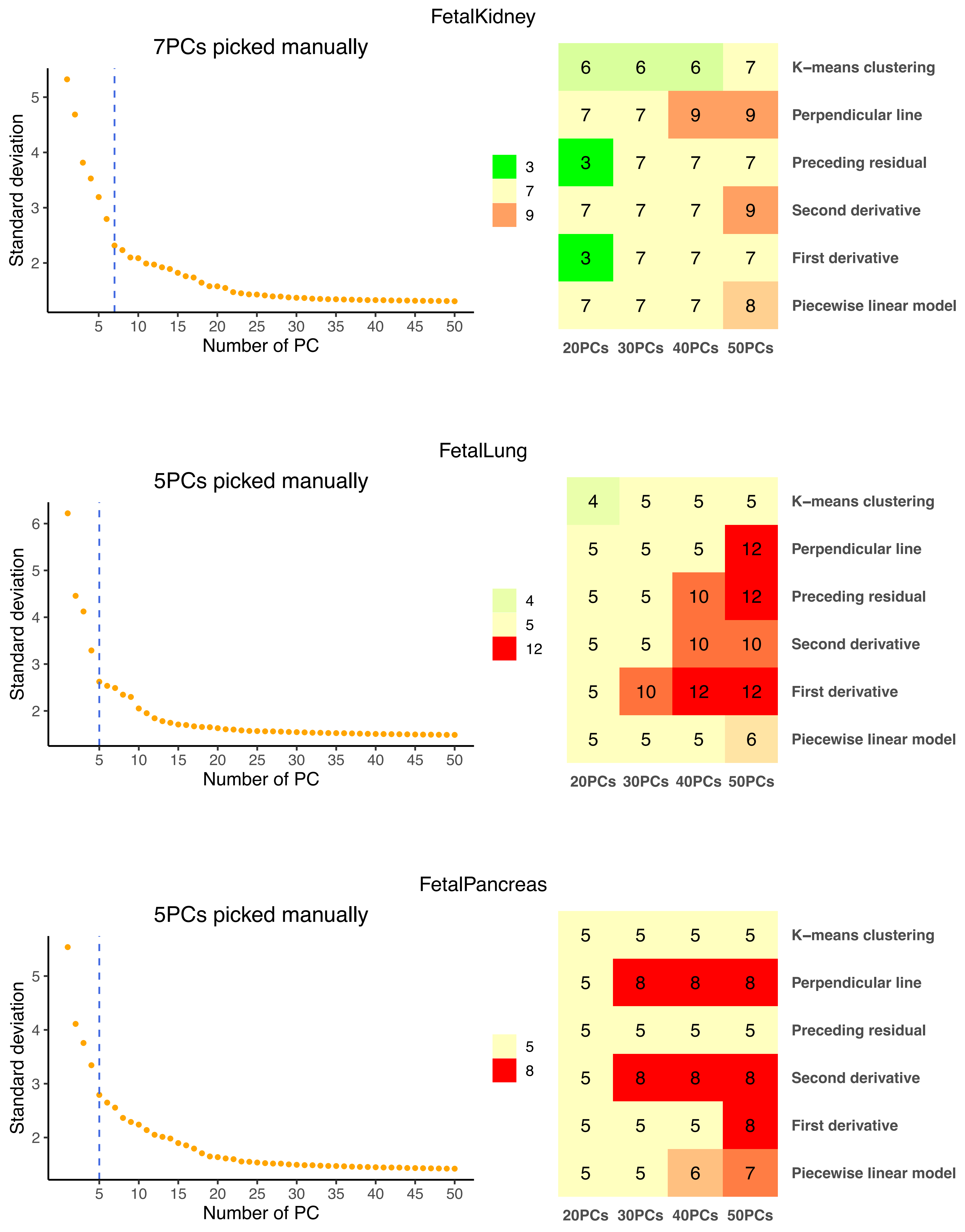

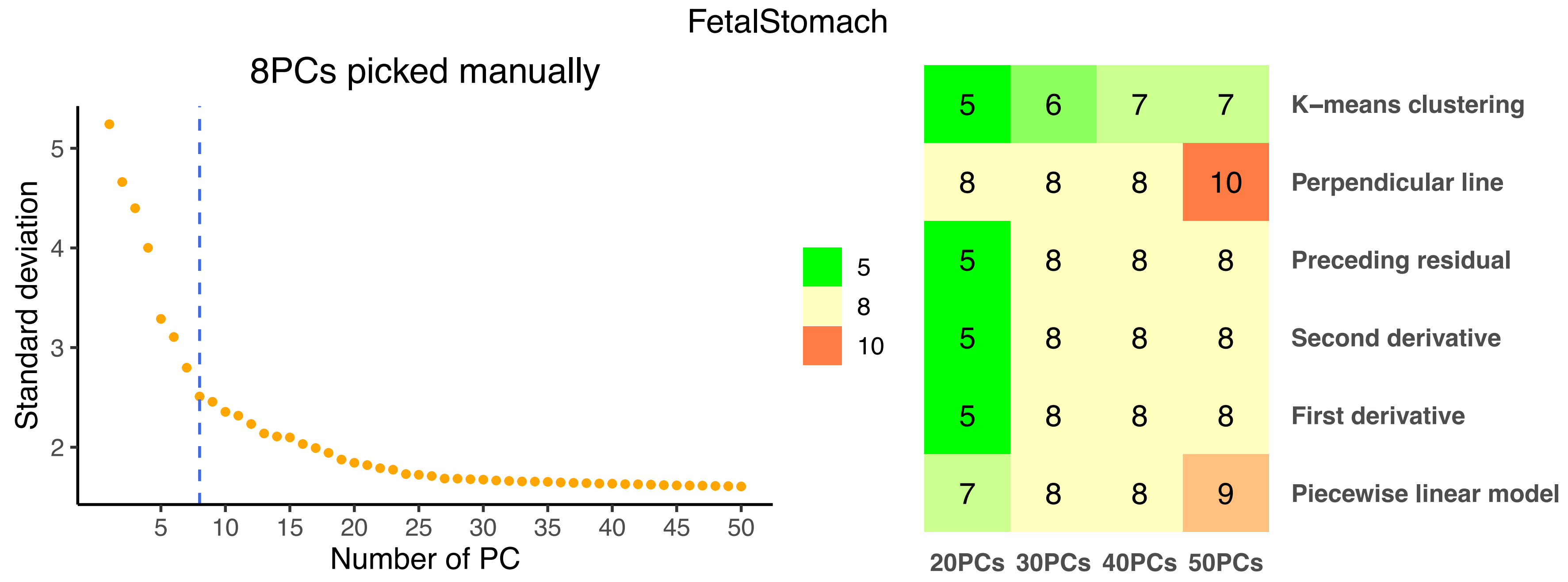

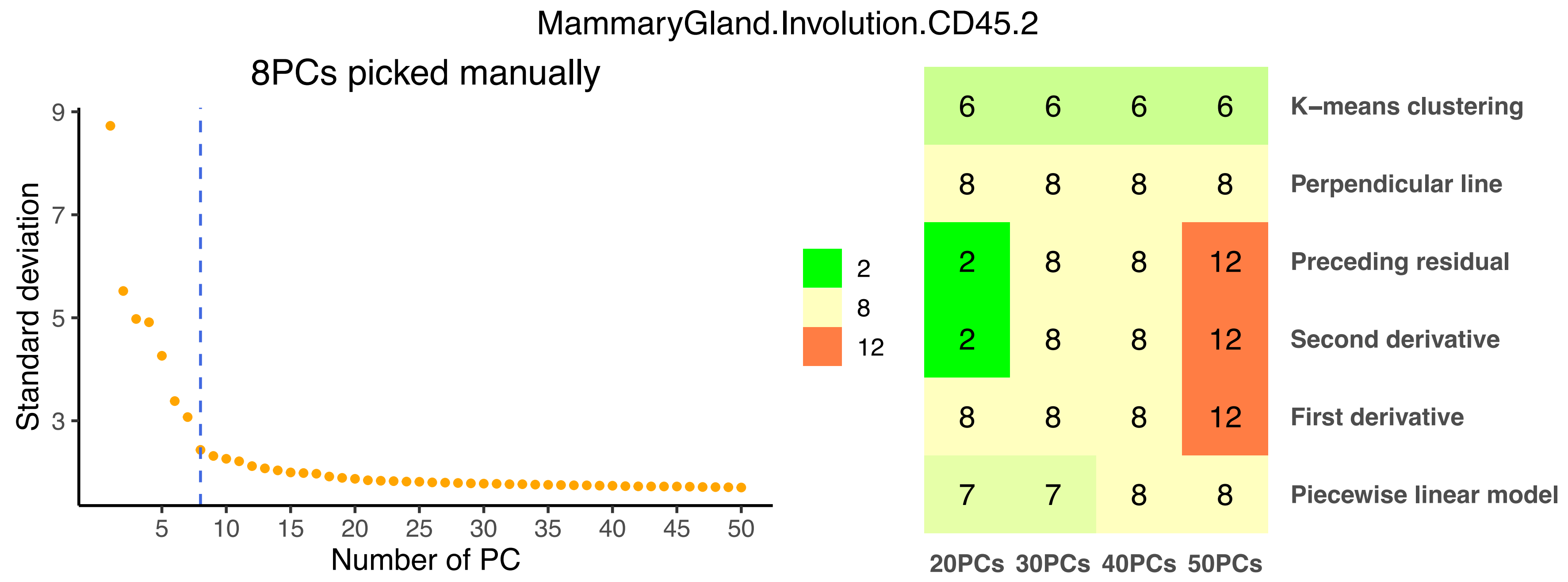

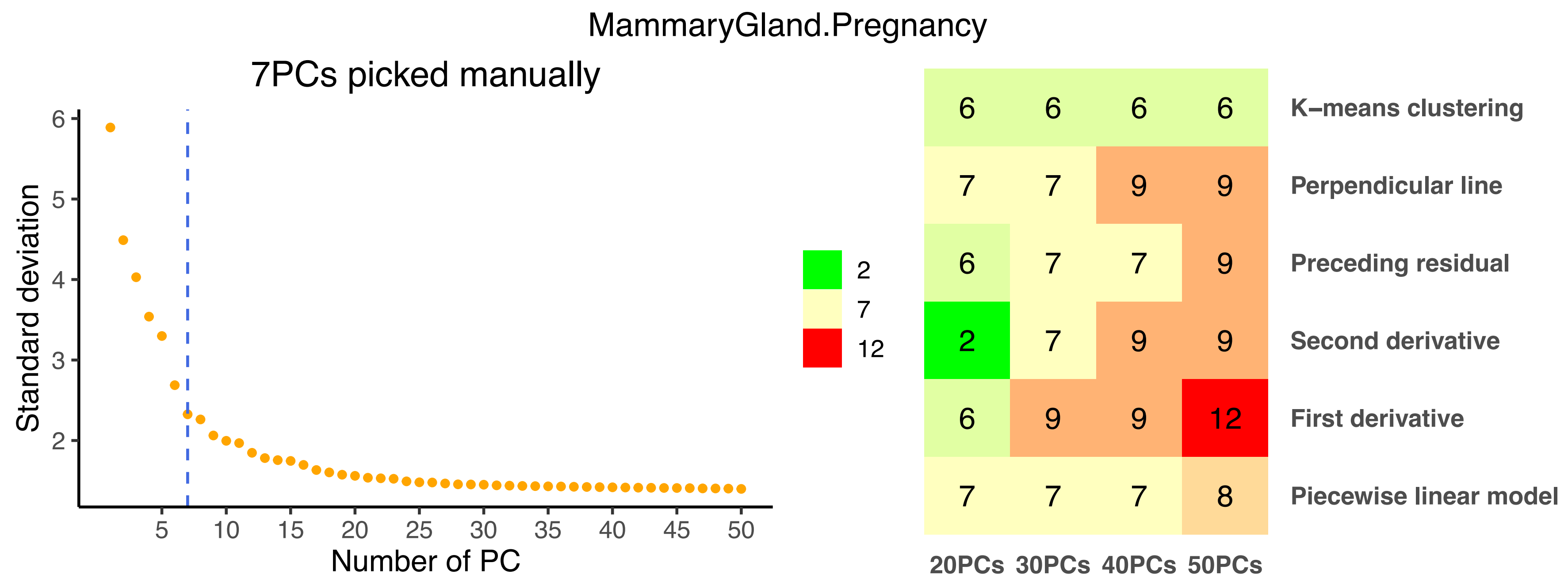

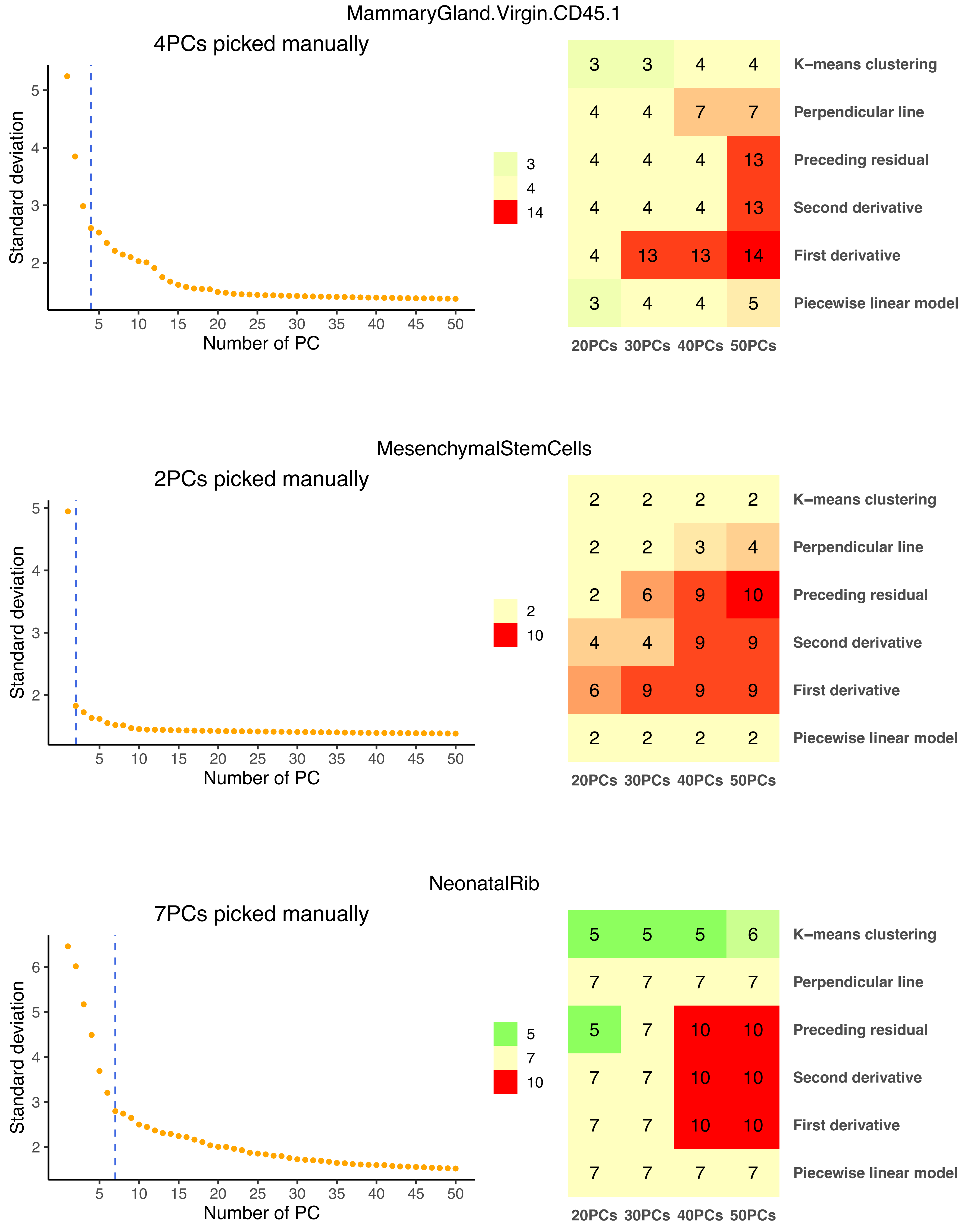

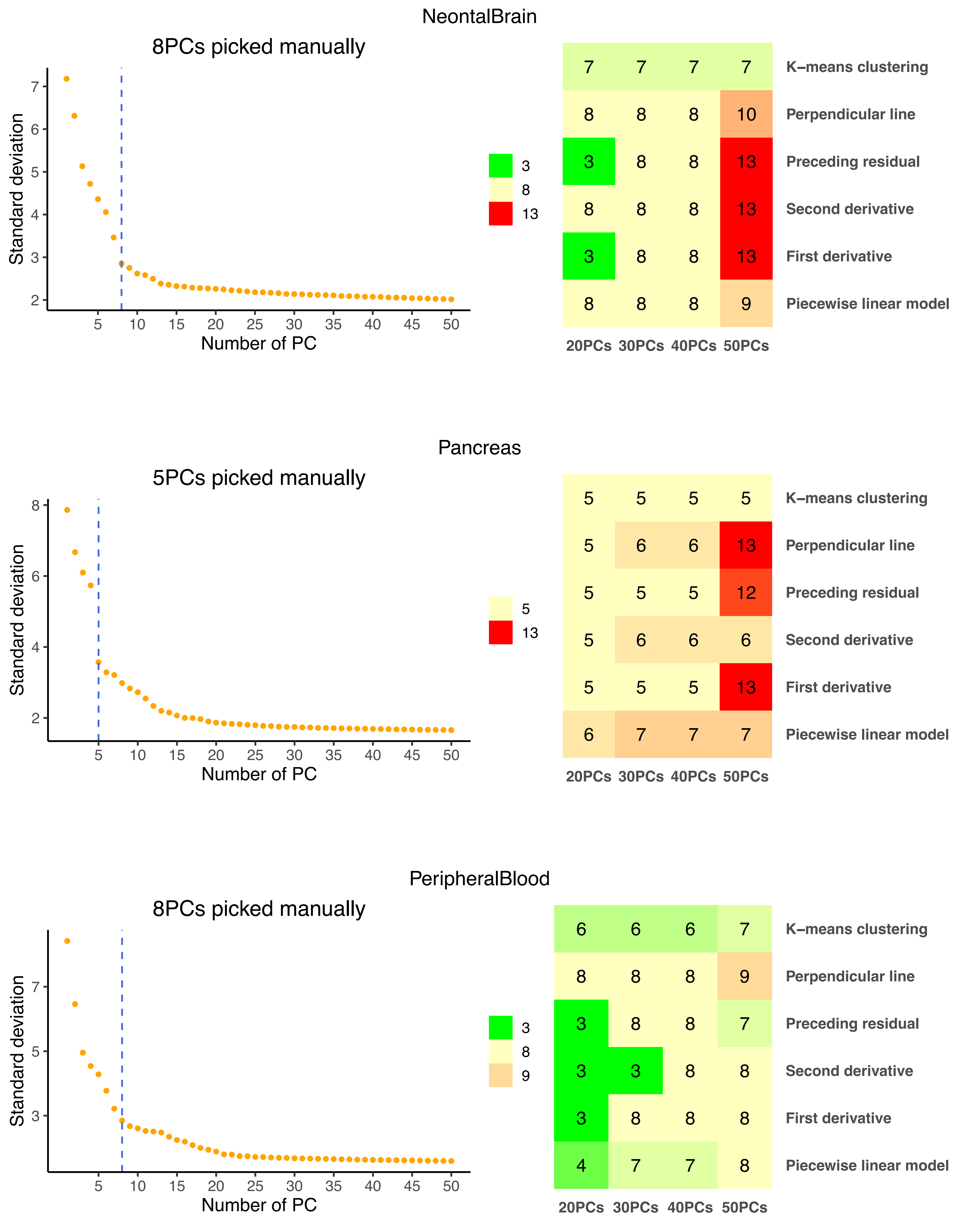

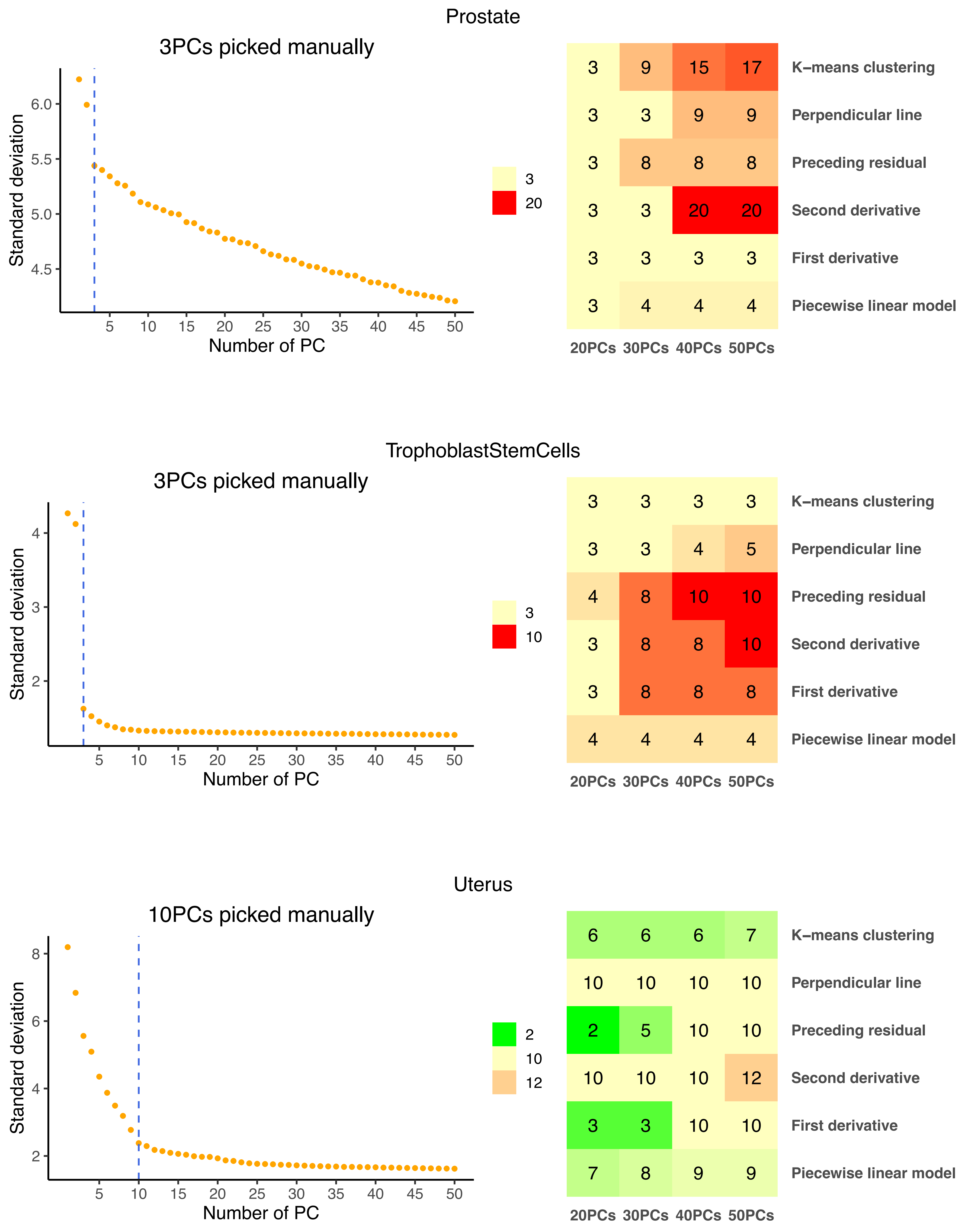

### Supplementary Figure 3

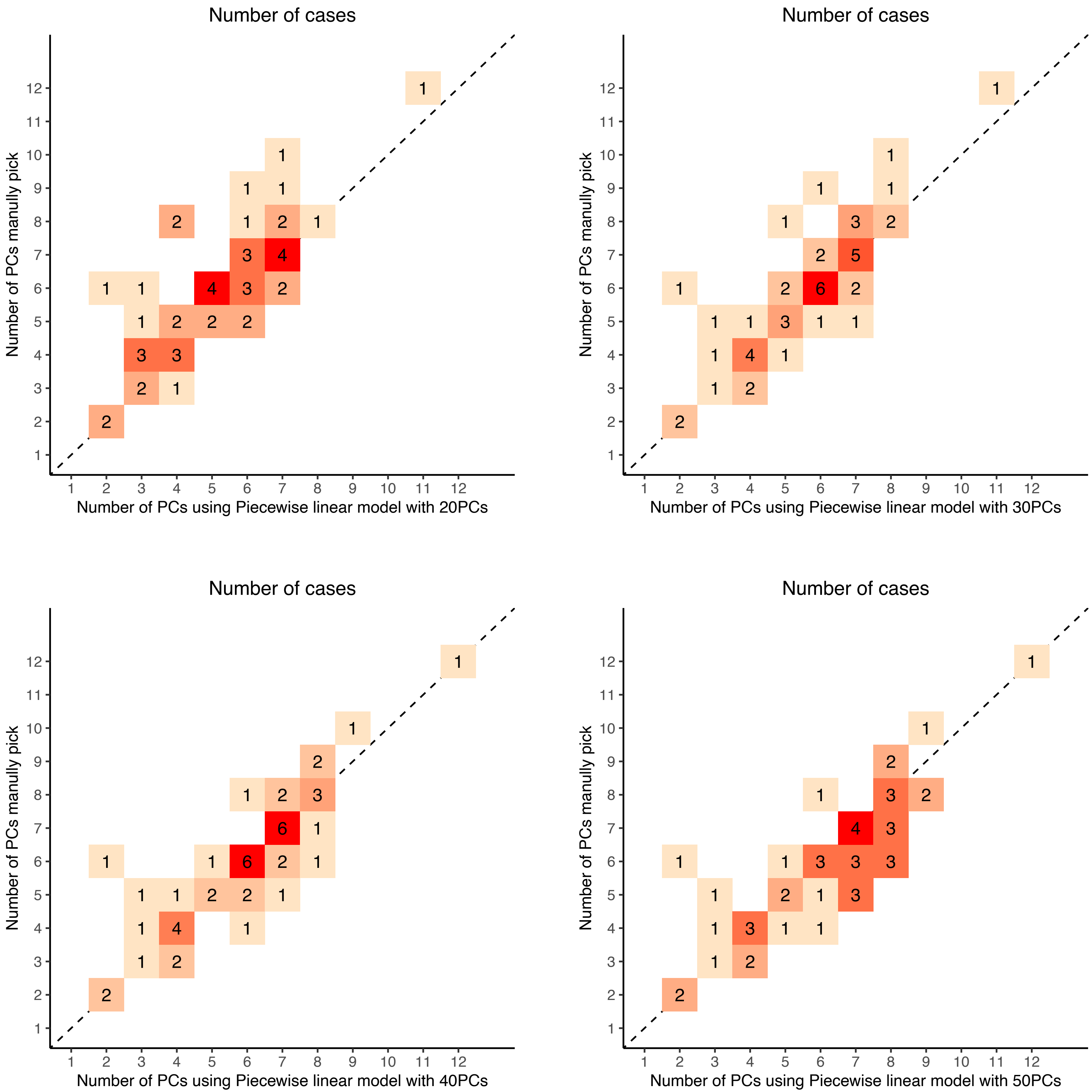

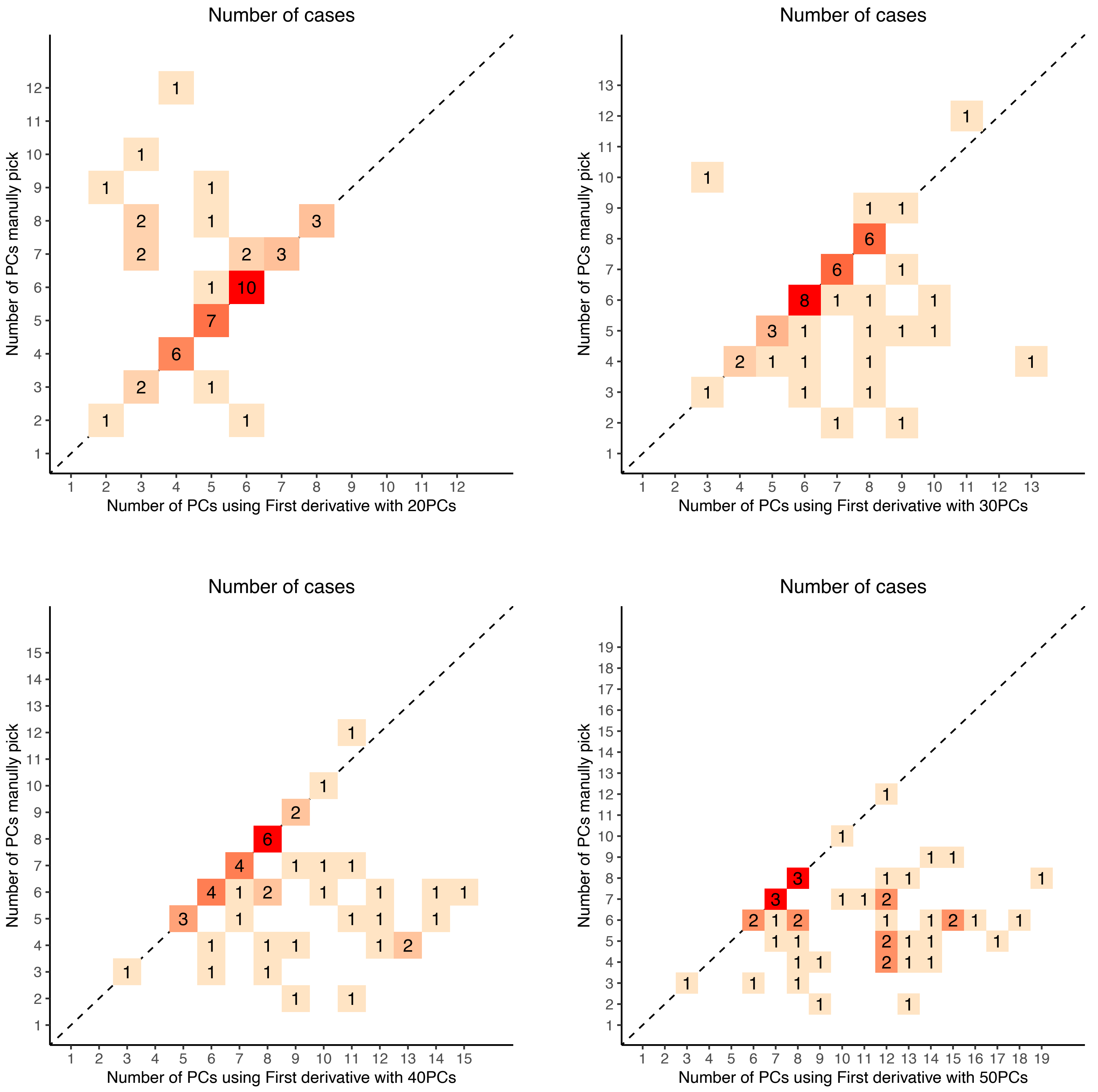

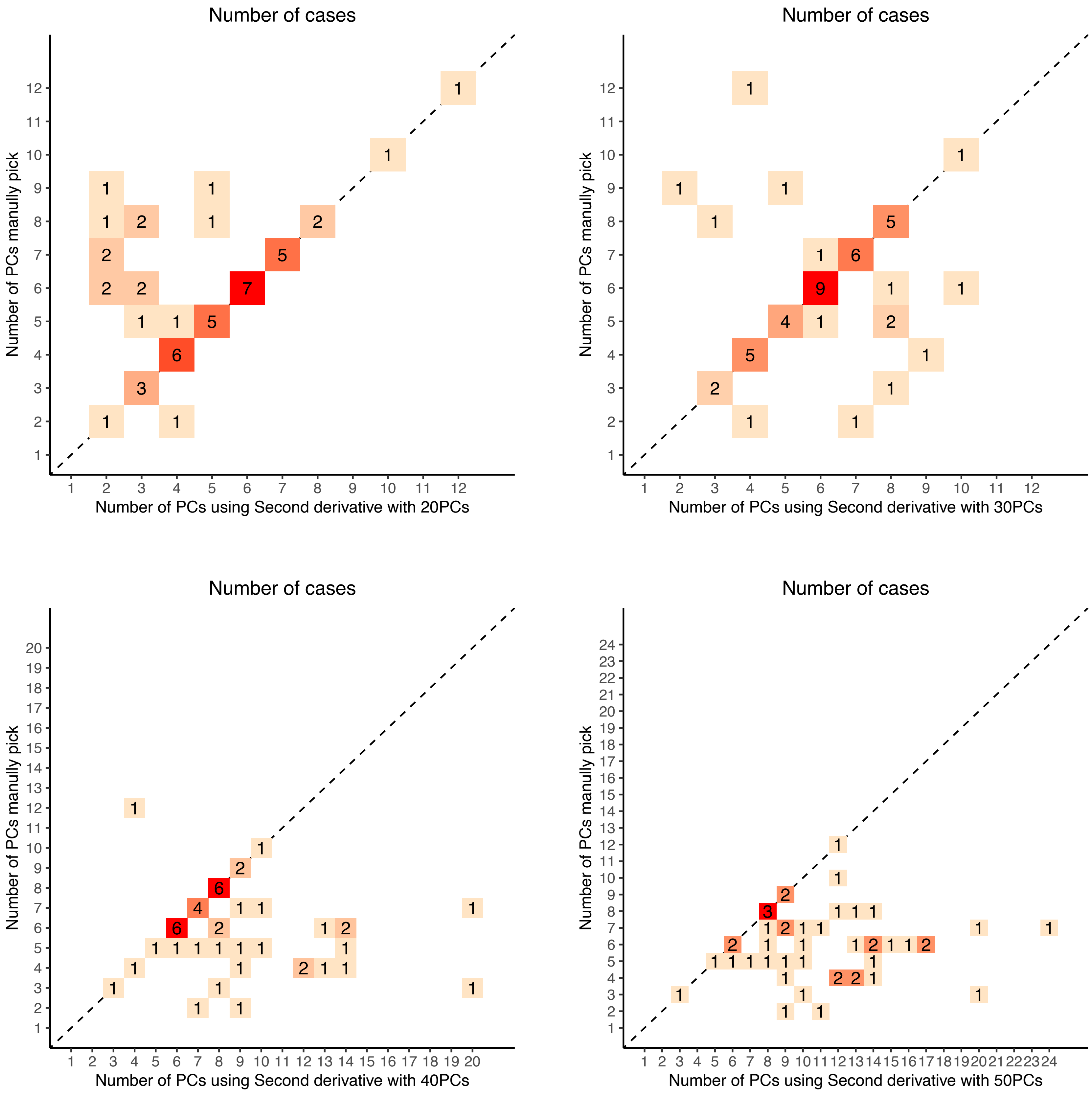

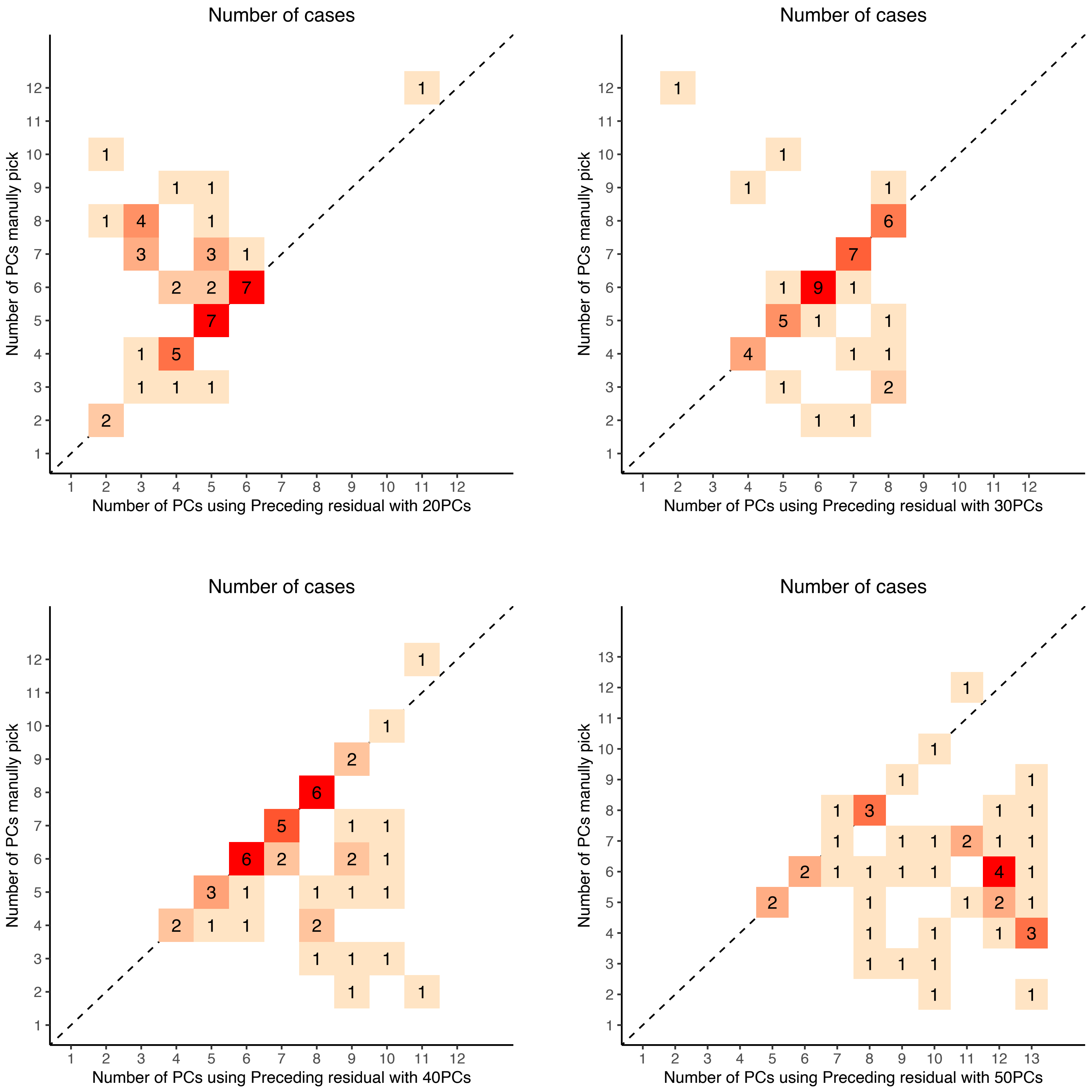

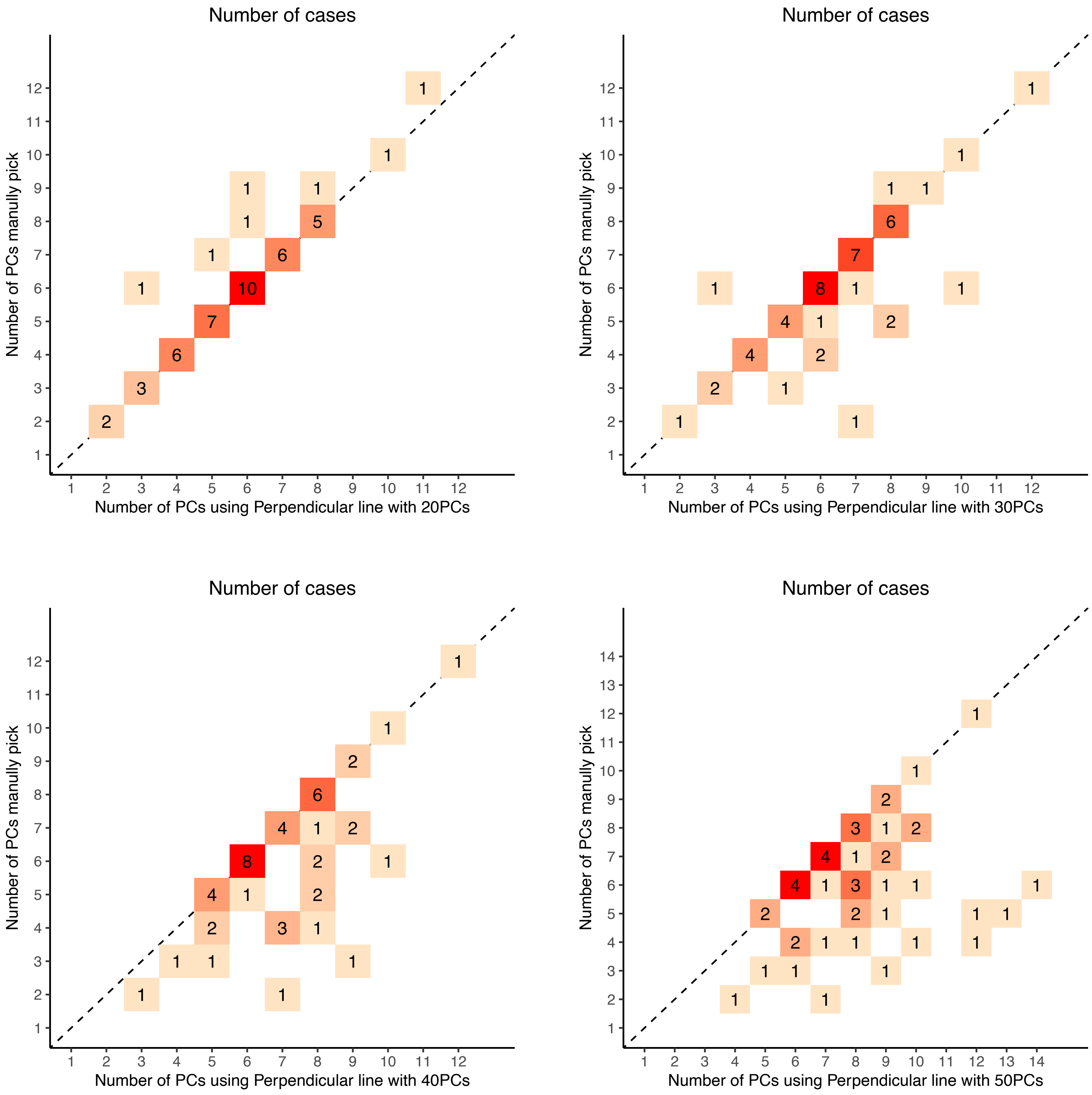

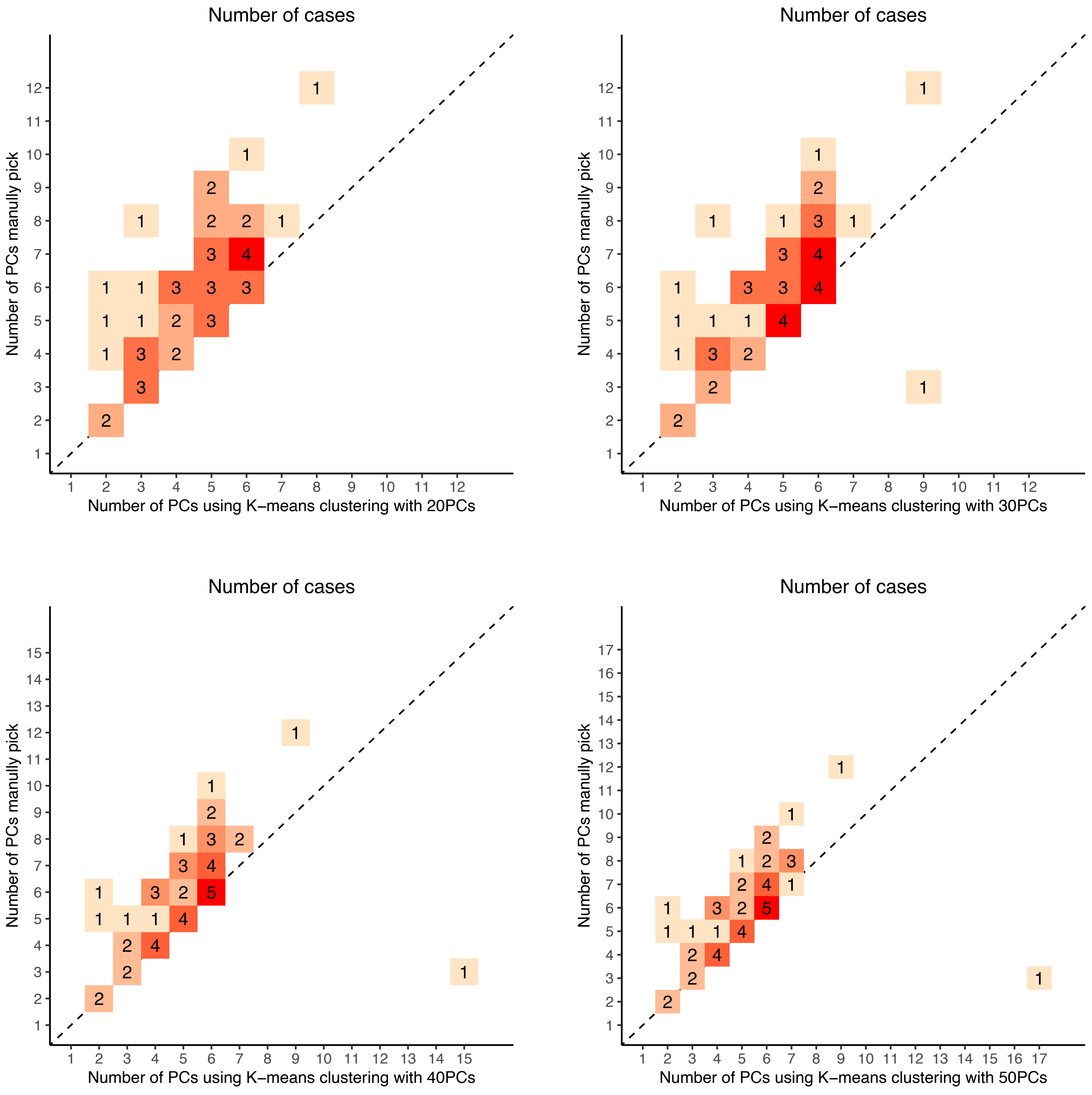
